# Organelle phenotyping and multi-dimensional microscopy identify C1q as a novel regulator of microglial function

**DOI:** 10.1101/2023.12.12.571151

**Authors:** Pooja S. Sakthivel, Lorenzo Scipioni, Josh Karam, Zahara Keulen, Mathew Blurton-Jones, Enrico Gratton, Aileen J. Anderson

## Abstract

Microglia, the immune cells of the central nervous system (CNS), are incredibly dynamic and heterogenous cells. While single cell RNA sequencing has become the conventional methodology for evaluating microglial state, transcriptomics do not provide insight into functional changes. Here, we propose a novel organelle phenotyping approach where we treat live human induced pluripotent stem cell-derived microglia (iMGL) with organelle dyes (mitochondria, lipids, lysosomes) and acquire data by live-cell spectral microscopy. Dimensionality reduction techniques and unbiased cluster identification allow for recognition of microglial subpopulations based in organelle function. We validate this methodology using lipopolysaccharide (LPS) and IL-10 treatment to polarize iMGL to an inflammatory” and “anti-inflammatory state, respectively, and then apply it to identify a novel regulator of iMGL function, complement protein C1q. C1q is traditionally known as the initiator of the complement cascade, but here we use organelle phenotyping to identify a role for C1q in regulating iMGL fatty acid storage and mitochondria membrane potential. Follow up evaluation of microglia with more traditional read outs of activation state confirm that C1q drives an increase in microglia pro-inflammatory cytokine production and migration, while suppressing microglial proliferation. These data together validate the use of a novel organelle phenotyping approach and enable better mechanism investigation of molecular regulators of microglial state, such as C1q.

## Introduction

As the immune cells of the central nervous system (CNS), microglia are constantly surveying the microenvironment via dynamic processes and protrusions to facilitate a rapid response to damage (Nimmerjahn et al., 2005). In response to neuropathological stimuli (such as CNS disease or injury), microglia respond within minutes by transitioning states via functional reprogramming (Hakim et al., 2021; Keren-Shaul et al., 2017). While it is well-understood that microglial state is no longer as binary as we once believed (Ransohoff, 2016), many microglial subpopulations that appear after disease/injury are associated with an inflammatory gene signature. These microglia release proinflammatory cytokines that are critical for the initial response to disease; however, prolonged cytokine release at the chronic stage of disease can inhibit neural repair and contribute to worsened pathology (Block et al., 2007; Polazzi & Contestabile, 2002; Zhang, 2011). Despite the notion that microglial inflammation heavily influences disease recovery, it remains unclear what molecular mechanisms regulate microglial inflammatory subpopulations in the healthy and diseased CNS.

One potential molecular regulator of interest is C1q, the initiator molecule of the complement cascade. While C1q is traditionally known for its role in the immune system, it has recently become recognized for novel functions as a single molecule within the CNS. For example, C1q regulates neurodevelopmental plasticity (Schafer et al., 2012; Stevens et al., 2007) and neurite outgrowth (Peterson et al., 2015). Here, we investigate a novel role for C1q in regulating microglial state and function. Indeed, C1q protein increases by 300-fold in the aged brain vs. control brain (Stephan et al., 2013), and further accumulates within the context of neurodegeneration (Yasojima et al., 1999); these data highlight the potential for C1q to influence CNS cells as a paracrine signal in these conditions. One previous study found C1q drives release of inflammatory cytokines (TNFa and IL6) in rat microglia primary cultures. In parallel with its traditional role within the complement cascade, we hypothesize that C1q is a molecular cue for regulating human microglial function and inflammation within the diseased CNS.

Critically, technological limitations have historically made it difficult to assess microglial responses to stimulation or disease at the resolution needed to discern complex heterogeneity. Recent advances in single-cell RNA sequencing (scRNAseq) allow for identification of numerous context-dependent microglial subpopulations, e.g., inflammatory and anti-inflammatory microglia (Michelucci et al., 2009), disease associated microglia (Keren-Shaul et al., 2017), interferon responsive microglia (Sala Frigerio et al., 2019), and white matter associated microglia (Safaiyan et al., 2021). Although scRNAseq identifies microglial heterogeneity and highlights pertinent signaling pathways, these naming conventions label subpopulations but are incapable of addressing the fundamental question of microglia function. Thus, caveats with scRNAseq that limit its applications include: (1) gene expression cannot predict the function of identified subpopulations; (2) transcriptomic changes often do not correlate with protein (Koussounadis et al., 2015); and (3) technical limitations do not allow for detection of lowly transcribed genes. Critically, a major gap in the field remains how to evaluate microglia functional heterogeneity at the single-cell level.

Here, we test a novel functional-based organelle phenotyping approach via multidimensional microscopy (Scipioni, 2023) to evaluate microglia organelles at a single-cell level. In contrast with traditional transcriptomic/proteomic techniques, organelle phenotyping provides quantifies real-time functional changes in energetic and anabolic metabolism. To achieve this, microglia are labeled with environment-sensitive dyes for mitochondria, lysosomes, and lipids, followed by data acquisition with spectral microscopy. This combination of organelle dyes and live microscopy allows for functional and morphological changes to be probed in individual, live cells following stimulation. This method has been validated using human induced pluripotent stem cell-derived microglia (iMGL) and quantification of cellular changes in response to the classical stimulants lipopolysaccharide (LPS) and IL-10. After validation, we apply this functional phenotyping approach to test the hypothesis that human purified C1q also influences microglial organelle functionality. These findings have been compared and validated by evaluation of classical transcriptional markers by qPCR. Overall, we identify that C1q influences microglial function via a myriad of outputs: organelles, gene expression, proliferation, and migration. These data suggest a novel function for C1q as a single ligand separate from its traditional role in the complement cascade, and highlight the capacity of organelle phenotyping to identify new molecules that regulate microglial states.

## Results

### Organelle phenotyping and multidimensional microscopy identify functional microglial subpopulations in an unbiased manner

Microglia undergo rapid transcriptomic changes in response to various molecular cues. However, changes in microglial organelles and their downstream impacts on cellular function (particularly in a single-cell, high throughput manner) are much less understood. Microglial state is associated with changes in metabolism, such as a mitochondrial shift towards glycolysis (Voloboueva et al., 2013) and an increase in acidic lysosomes (Majumdar et al., 2007) and lipids (Button et al., 2014). These readouts are much more indicative of cell function when compared to transcriptomics because organelle changes are direct readouts of cell metabolism. We therefore hypothesize that organelle phenotyping and multidimensional microscopy can identify microglial subpopulations by probing changes in microglial organelles as an alternative output to study activation.

As a validation for this methodology, iMGL either remain untreated as a negative control, or are treated with LPS or IL-10 for 24 hours. Although we acknowledge microglia are non-binary cells, the extreme ends of microglial heterogeneity can be described via the “inflammatory” and “anti-inflammatory” nomenclature. Because LPS and IL-10 have traditionally been used to polarize microglia to these states, we use these stimulants as positive controls to investigate whether this methodology could identify microglial subpopulations in an unbiased manner. Because a previous study has suggested a role for C1q in regulating cytokine release in rodent microglia (Färber et al., 2009), iMGL are also treated with purified C1q to test the hypothesis that C1q similarly regulates human microglial state and function. We predict C1q polarizes microglia towards an inflammatory phenotype and thus expect to observe a similar but distinct pattern when compared to LPS-treated cells. In addition to stimulants, iMGL are labeled with the following environment-sensitive dyes to observe changes in organelles: Rhodamine123 (mitochondrial membrane potential), lysotracker green (lysosomal pH), nile red (lipid droplet polarity), and hoechst (DNA) **(Figure 1A-D)**. Data is acquired using live-cell spectral microscopy, allowing for quantification of organelle characteristics with single-cell resolution.

**Figure 1:**
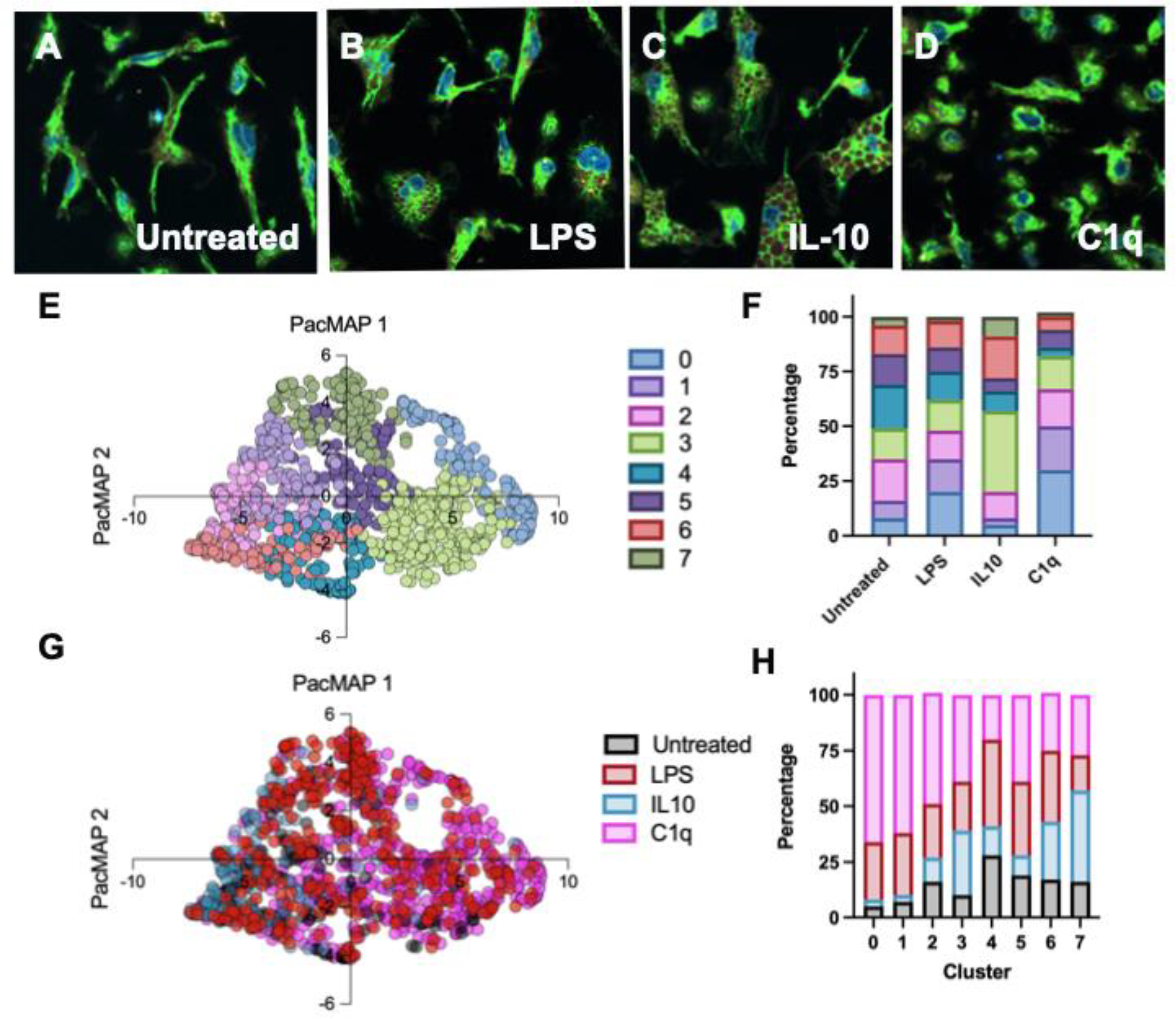
C1q triggers changes in iMGL organelles and morphology, similar to LPS. (A-D) iMGL remain untreated as a control (A) or are treated with LPS (B), IL-10 (C), or C1q (D). Cells are labeled with lysotracker green (lysosomes, cyan), Nile red (lipids, yellow), Rhodamine 123 (mitochondria, white), and hoechst (DNA, magenta). Live cells are imaged 24 hours following treatment. (E-H) Data acquisition is followed by PacMAP dimensionality reduction and cluster identification, revealing 8 distinct cluster that are associated with treatments (E, G). Cluster composition by treatment (F) and treatment composition by cluster (H) show cluster analysis is able to recognize heterogeneity and functional changes driven by treatment in an unbiased manner.

Data acquisition is followed by PacMAP dimensionality reduction and unbiased subpopulation identification. The data reveals 8 distinct microglial subpopulations **(Figure 1E)**; cluster distribution by treatment shows that certain clusters are predominantly rooted in one treatment group **(Figure 1F)**. PacMAP can alternatively be visualized by treatment **(Figure 1G)**, highlighting how treatments differentially influence cells. We observe untreated cells cluster separate from the treated cells. Moreover, IL10 cells separate out but LPS and C1q treated cells seem to be overlapped, suggesting LPS and C1q-treatments drive iMGL to behave similarly. We can additionally quantify treatment distribution by cluster **(Figure 1H)**, thereby enabling correlations to be drawn between clusters/treatments and allowing for downstream analysis comparing treatment groups. Clusters 0, 1, 2, and 5 are associated predominantly with C1q and LPS treated iMGL. Because four out of seven clusters share identification between C1q and LPS-treated cells, this data supports the hypothesis that C1q drives changes in organelle function that are consistent with a proinflammatory phenotype. Clusters 3 and 7 are largely made up of IL-10 treated cells, whereas cluster 4 shows untreated cells. Lastly, cluster 6 is comprised of all four treatment groups suggesting it may represent a baseline homeostatic or trophic subpopulation present despite treatment.

As we predicted, organelle phenotyping identifies changes in microglia in response to stimulation. PacMAP followed by unbiased cluster identification finds distinct cell subpopulations in an unbiased manner. Therefore, organelle phenotyping is a useful methodology for characterizing microglial states based on function and offers an alternative method of quantification for studying microglial state.

### C1q drives complex changes in iMGL morphology and organelle function

These resulting subpopulations can be analyzed to identify which characteristics are driving cluster separation, thereby providing unbiased insight into which characteristics are most different across cell treatments and clusters.

Morphology is one the first prominent changes observed in microglia within the diseased CNS (Nimmerjahn et al., 2005). While homeostatic microglia are typically branched, activated microglia adopt larger cell volume and decreased branching complexity. As expected, iMGL increase cell area and decrease cell solidity (a readout of how not branched the cell is) in response to all treatments. Morphology of C1q-treated cells remains significantly different when compared to LPS or IL-10 treated cells, suggesting C1q may be driving a distinct phenotype separate from how classical stimulants influence microglial state **(Figure 2A-B)**. Overall, all three treatments drive the same trend of a larger, more amoeboid morphology, but the extent of morphological changes across treatments remain distinct. These data suggest that, while all stimulants are triggering microglial activation, the stimulants may be functioning through different mechanisms and thereby have discrete impacts on cell function.

**Figure 2:**
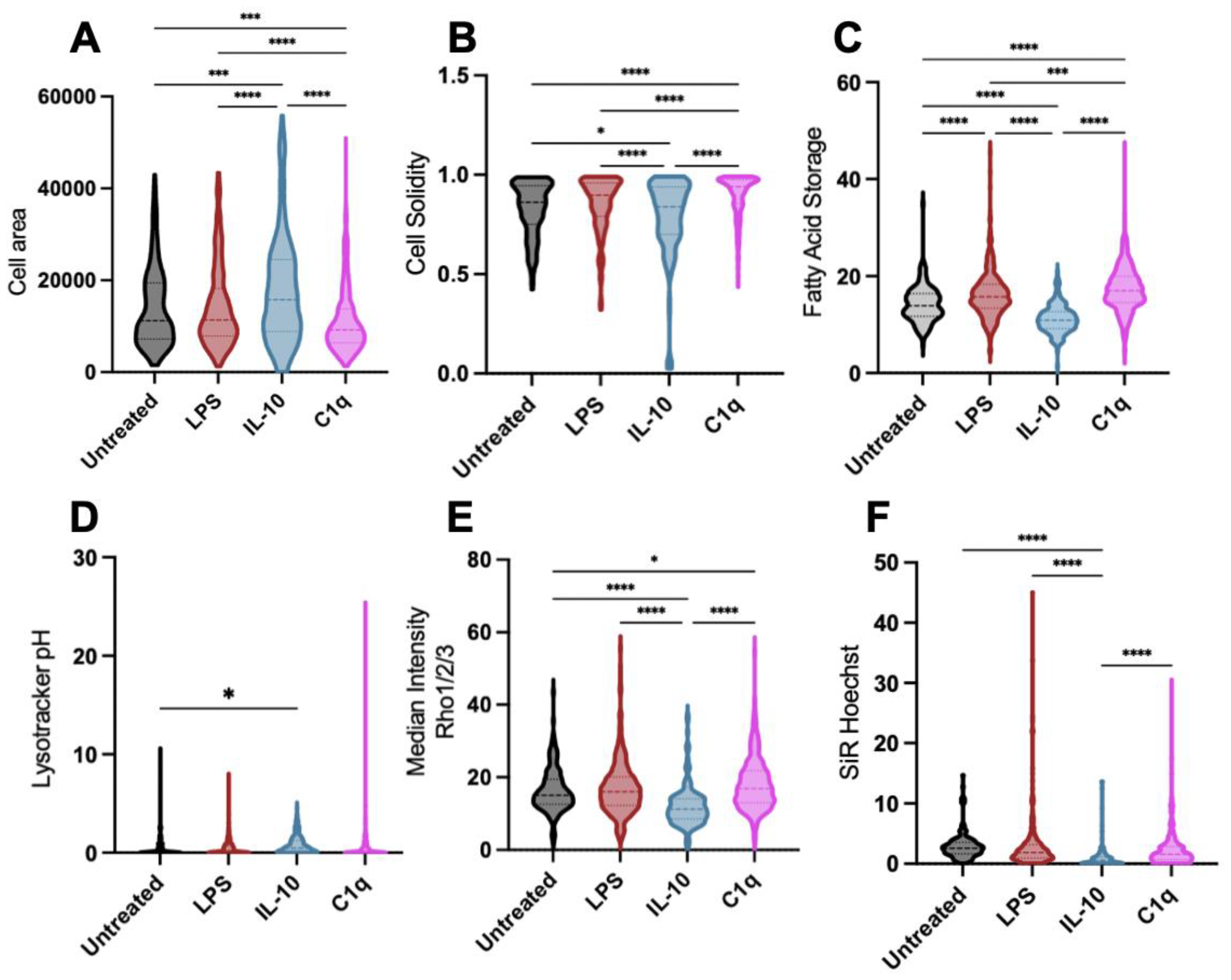
Functional changes in microglial organelles drive cluster separation. Subpopulation identification in an unbiased manner is largely due to changes in microglial morphology (A-B), lipids (C), lysosome acidity (D), mitochondria membrane potential (E), and DNA condensation (F). Statistical analysis by one-way ANOVA (p≤0.05) and Tukey’s t-test as indicated. *p≤0.05, **p≤0.01, ***p≤0.001, ****p≤0.0001.

Increased fat storage accumulation and accumulation of lipid droplets are associated with inflammation and activation in immune cells (D’Avila et al., 2008). Indeed, LPS drives an increase in fatty acid storage **(Figure 2C)** as has been previously noted in the literature for murine microglia (Khatchadourian et al., 2012). IL-10 treatment conversely drives a slight decrease in fatty acid storage in comparison to untreated microglia, suggesting fat storage is associated with proinflammatory microglia but not anti-inflammatory microglia. Interestingly, microglia treated with C1q display an increase in fatty acid storage that is comparable to the LPS treatment. These data show that both LPS and C1q are extracellular cues that are sufficient to induce increased fatty acid storage, suggesting proinflammatory cells display increased fat storage in a way homeostatic and anti-inflammatory microglia do not.

As the primary phagocytes within the CNS, microglia and their lysosomes are often indicative of phagocytosis and digestion. Lysosomal pH is a key characteristic in separating out IL-10 treated iMGL from other treatments **(Figure 2D)**. We show IL-10 triggers an increase in lysosome acidity, consistent with previous findings showing IL-10 treatments increase phagocytic capacity of microglia (Yi et al., 2020). In contrast, neither LPS not C1q treatments alter lysosomal pH. These data suggest anti-inflammatory microglia have more acidic lysosomes, likely due to increased phagocytosis or digestion demands observed in response to IL-10 treatment.

We have also observed changes in mitochondrial membrane potential across treatments **(Figure 2E)**. LPS does not change Rho123 intensity, but IL-10 drives a significant decrease and C1q drives a significant increase. Interestingly, this is one organelle phenotype where LPS and C1q display separate trends, suggesting C1q likely promotes microglial oxidative phosphorylation and ATP production in a manner that is distinct from LPS. These differing trends across all treatments highlight the variable changes in energy dynamics and the need for a functional output like mitochondria membrane potential.

We lastly identify intensity of SiR-hoechst (DNA stain) as a characteristic that separates out the identified subpopulations **(Figure 2F)**. We observe a significant decrease in SiR-hoechst intensity following IL-10 treatment, suggesting chromatin unfolding. This is validation of IL-10 and its influence on epigenetic modifications that has previously been noted in the literature (Rajbhandari et al., 2018). While LPS and C1q show a trend towards increased SiR-hoechst intensity which would suggest higher chromatin compaction, the trend is not statistically significant for either treatment. Chromatin changes following LPS treatments have also been noted in the literature in other cell types (Shen et al., 2016). Overall, DNA/chromatin is a meaningful output to observe when considering how epigenetics could be altering microglial state.

In summary, we demonstrate organelle phenotyping can simultaneously investigate multiple organelle outputs associated with microglial state in an unbiased manner. Using this tool, we show C1q induces microglial organelles to exhibit characteristics similar to a proinflammatory state but distinct from the effects of LPS.

### C1q drives an inflammatory gene signature in iMGL, as predicted by the organelle changes observed with multidimensional microscopy

Because LPS and C1q treated iMGL show similar trends in organelle phenotyping, we predict that LPS and C1q treated iMGL would also show similar trends in RNA production, a readout that is more historically used to evaluate microglial state. We thus use qPCR as a more traditional evaluation of gene expression to assess microglial state and quantify 3 representative inflammatory, anti-inflammatory, and homeostatic genes. To additionally investigate a dose-response, iMGL are treated with three physiologically relevant concentrations of C1q: as produced by neutrophils [0.1 nM], macrophages [1 nM], and is present in the blood/serum [200 nM] (Hooshmand et al., 2017). After a 24 hour C1q treatment, iMGL are lysed and mRNA is isolated for qPCR to investigate changes in transcription. Consistent with LPS, C1q also triggers an upregulation of inflammatory genes (CCL2, IL1b, IL6) in a dose-dependent manner **(Figure 3A)**. These data confirm that C1q is sufficient to drive an inflammatory gene profile. Thus, the microglial phenotypes identified with multidimensional microscopy (**Figure 1**) are correlated with changes in gene expression that are associated with an inflammatory state.

**Figure 3:**
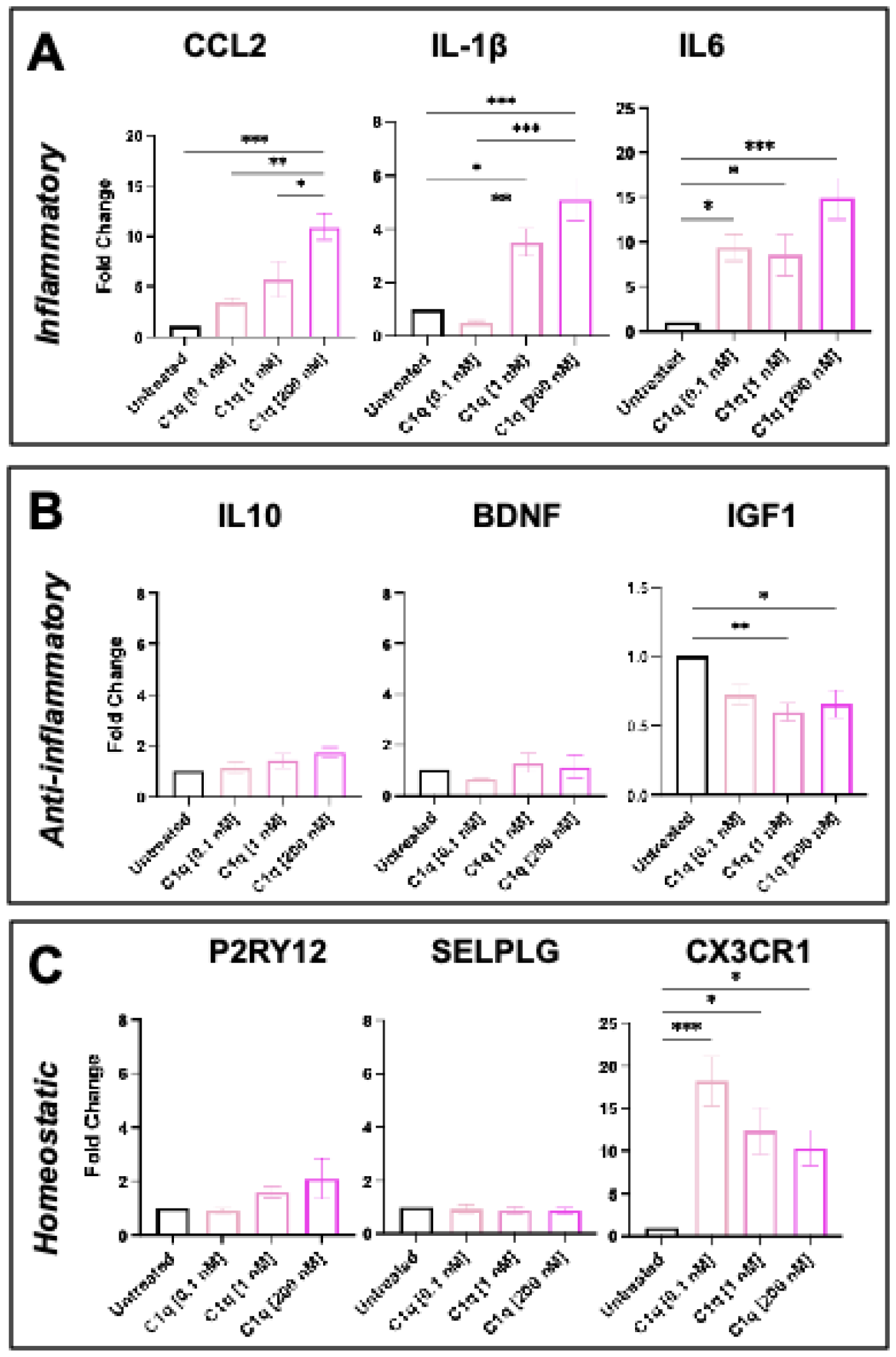
Paracrine C1q drives an inflammation-associated transcriptional signature. iMGL are treated with C1q at three physiologically relevant concentrations: 0.1 nM, 1 nM, and 200 nM. Cells are lysed following 24 hour treatment and mRNA is collected to investigate (A) inflammatory, (B) anti-inflammatory, and (C) homeostatic gene expression. Data shows mean ± SEM normalized to untreated controls. Statistical analysis by one-way ANOVA (p≤0.05) and Tukey’s t-test as indicated. *p≤0.05, **p≤0.01, ***p≤0.001, ****p≤0.0001. n=3-4 biological replicates.

We additionally probe whether C1q changes the expression of representative anti-inflammatory or homeostatic genes. C1q has no effect on anti-inflammatory genes (IL-10 and BDNF), but decreases the anti-inflammatory marker IL6 **(Figure 3B)**. Because IL6 can stimulate C1q production (Faust & Loos, 2002), this may be a mechanism used to inhibit C1q production and thus ultimately exit an inflammatory phenotype. Lastly, C1q does not influence homeostatic genes P2RY12 and SELPLG. However, C1q does drive an increase in CX3CR1 **(Figure 3C)**. While CX3CR1 has historically been thought of as a homeostatic marker, CX3CR1 has recently been implicated in inflammatory processes as well (Freria et al., 2017; Ho et al., 2020). Interestingly, CX3CR1 additionally has a known role in regulating phagocytosis (Zabel et al., 2016), thus, this may be a mechanism underlying C1q-driven phagocytosis, as aligned with C1q’s traditional function (Fraser et al., 2010). Overall, these data demonstrate that paracrine C1q influences microglia to adopt an inflammatory gene signature.

### C1q triggers iMGL chemotaxis and phagocytosis, while attenuating iMGL proliferation

Microglia rapidly respond to changes in homeostasis by migrating towards regions of disease, proliferating, and/or inducing apoptosis. We therefore lastly evaluate iMGL responses to C1q via quantitative assays that characterize microglial functions distinct from their transcriptomic signature, providing us with a better understanding of how C1q influences microglial behavior and not just gene signature.

Because microglia are highly dynamic and motile cells within the CNS, we test if C1q can serve as a chemotactic cue for iMGL using a transwell migration assay. We find that iMGL exhibit increased migration towards all doses of C1q. While all doses trigger chemotaxis, C1q[1 nM] interestingly induces increased migration when compared to C1q[0.1 nM] and C1q[200 nM], highlighting the dose-dependent influence C1q on iMGL migration **(Figure 4A)**.

**Figure 4:**
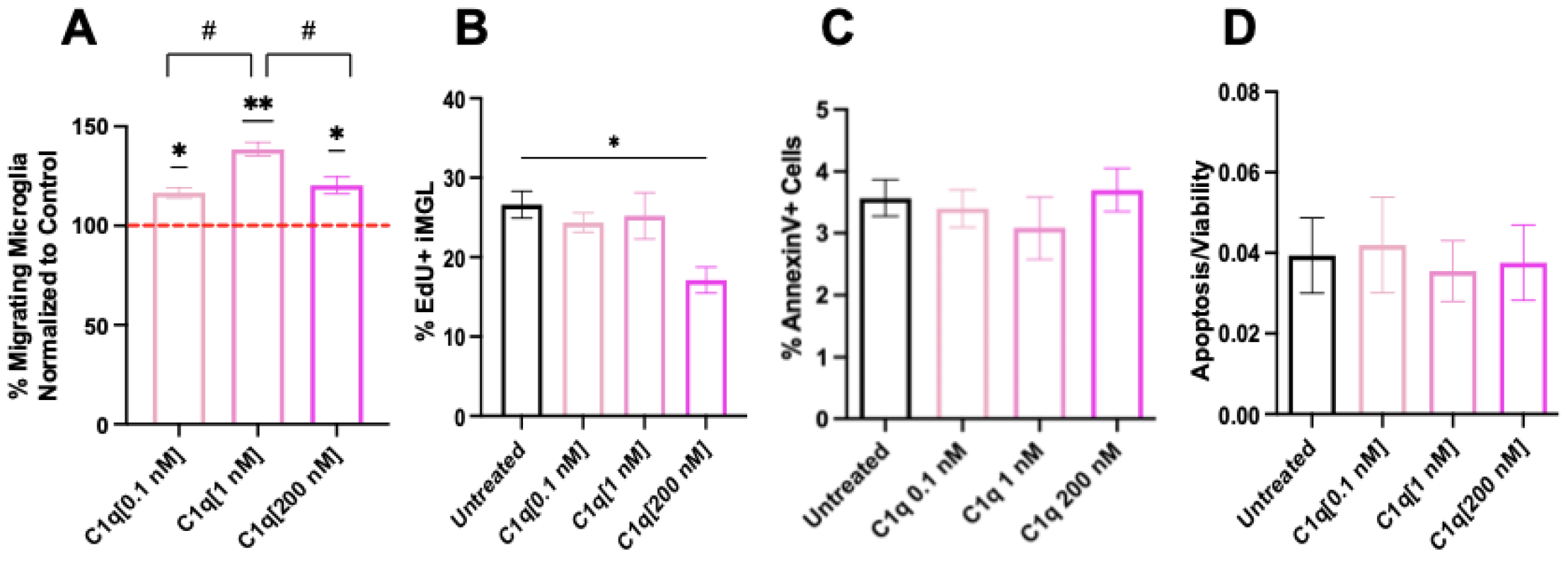
C1q[200 nM] induces a decrease in microglial baseline proliferation and decreased cell death. (A) C1q[0.1 nM], C1q[1 nM], and C1q[200 nM] all induce iMGL chemotaxis in transwell migration assays. C1q[1 nM] induces an increased level of migration, in comparison to low and high C1q doses. Statistical analysis using one sample t-test (*p≤0.05, **p≤0.01) for comparison with untreated control and using one-way ANOVA (p≤0.05) followed by Tukey’s post-hoc t-test as indicated (#p≤0.05) for comparison between conditions. n=3 biological replicates. (B-D) iMGL are treated with C1q[0.1 nM], C1q [1 nM], or C1q[200 nM] for 24 hours and processed to quantify proliferation and cell death. (B) C1q[200 nM] drives a significant decrease in the percentage of EdU+ cells in comparison to untreated iMGL. (C,D) C1q treatment does not influence microglia cell death, as quantified by AnnexinV staining and flow cytometry (C) and Apolive Glow assay (D). Statistical analysis by one-way ANOVA (p<0.0001) and Tukey’s multiple comparisons test as indicated. **p≤0.01, ****p≤0.0001. n = 3-4 biological replicates.

We also assess cell proliferation via EdU incorporation and apoptosis via AnnexinV expression following 24 hours of C1q treatment. These data demonstrate that approximately 25% of iMGL proliferate at baseline; however, this is significantly reduced in the presence of C1q[200 nM] **(Figure 4B)**. This data also shows that C1q has no influence on apoptosis (AnnexinV+ cells) **(Figure 4C)**. These findings have been independently replicated using the ApoliveGlo viability assay that provides a luminescent readout of Caspase 3/7 activity as an alternative method to quantify apoptosis **(Figure 4D)**. Overall, we find that C1q induces iMGL migration at all doses and attenuates iMGL proliferation at C1q[200 nM] only, but has no influence on iMGL apoptosis or overall viability. These data show that C1q influences complex changes in microglial cell function by controlling migration and cell proliferation.

## Discussion

Early studies in the mid-1970s first coined the term “microglial activation” when observing the striking difference in microglial morphology following brain damage (Tremblay et al., 2015). It is now understood that “microglial activation” is not black and white and better tools are needed to evaluate these complex cells (Paolicelli et al., 2022). Here, we hypothesize that changes in organelles are a well-suited methodology to investigate the wide spectrum of microglial polarization. We utilize a novel multidimensional spectral microscopy technique to quantify changes in microglial organelles and investigate microglial subpopulations in an unbiased, functionality-based manner. We demonstrate the capacity to quantify changes in microglia mitochondria, lipids, and lysosomes, as well as unbiased subpopulation clustering. We first test the classical stimulants LPS and IL-10 to validate this functional phenotyping approach. We show that LPS increases lipid number, whereas IL-10 increases lysosomal pH and decreases mitochondria membrane potential. These data suggest inflammatory cells are likely to increase fat storage, whereas anti-inflammatory microglia increase lysosomal digestion and decrease energy production. We next apply this method to investigate the hypothesis that C1q influences microglial state. Indeed, we show that C1q increases fatty acid storage in iMGL as LPS does, consistent with an inflammatory phenotype. However, unlike LPS, C1q also upregulates mitochondria membrane potential, suggesting C1q drives an iMGL phenotype that is distinct from that of LPS. This critically suggests that C1q treatment increases energy production in microglia and provides a functional understanding of how C1q is modulating microglial state. These data together identify C1q as a novel and unique molecular regulator of microglial state/function.

The complement cascade is the immune system’s first line of defense against pathogens, functioning by initiating a series of enzymatic reactions, recruiting inflammatory cells, tagging pathogens for removal, and triggering cell lysis. C1q, the recognition molecule of the classical complement cascade, has a well-defined role in recognizing antigens and pathogens, initiating autocatalytic activation of the complement cascade, and tagging debris for clearance by phagocytic immune cells. In addition to its traditional roles in the immune system, C1q as a single molecule has recently become recognized for novel functions within the CNS. Indeed, C1q induces ERK, mitogen-activated protein kinases (MAPK), and Akt signaling pathways (Agostinis et al., 2010; Benavente et al., 2020; Lee et al., 2018), suggesting it could play a more complex role than just a phagocytic tag. In support of this notion, C1q mediates neurodevelopmental plasticity by tagging low-activity presynaptic terminals for elimination by microglia (Schafer et al., 2012; Stevens et al., 2007), modulates axon growth by suppressing myelin-associated glycoprotein growth inhibitory signaling (Peterson et al., 2015), and serves as a chemotactic cue to initiate neural stem cell migration (Benavente et al., 2020). In parallel to our study, extrinsic C1q stimulates inflammatory cytokine (TNF-α, IL-6 and nitric oxide) release from rodent microglia (Färber et al., 2009), suggesting the role of C1q as an inflammatory cue is conserved across species.

C1q is present within the healthy and intact CNS, but is highly upregulated following aging (Stephan et al., 2013), injury (Figley et al., 2014), ischemia (Schäfer et al., 2000), Alzheimer’s disease (Chatterjee et al., 2023), Parkinson’s disease (Depboylu et al., 2011), and numerous other neurodegeneration conditions. These data suggest that C1q is likely to play a broad role in modulation of CNS inflammation in the context of disease and injury. In this regard, treatment of primary rat microglia cultures with exogenous C1q increases proinflammatory cytokine and nitric oxide release (Färber et al., 2009). Here, we show that C1q similarly increases human iMGL transcription of proinflammatory markers in a dose-dependent manner. Therefore, C1q in the diseased CNS is likely to be modulating microglial inflammation associated with aging, neurodegenerative disease, and injury.

Overall, the data in this manuscript therefore identifies a newfound mechanism for which microglial setpoints are initiated and maintained through C1q, a molecule that has biological relevance across a range of neurodegenerative disorders. While this exogenous (paracrine) C1q clearly influences microglial inflammation, autocrine C1q has not yet been investigated as a distinct mechanism. Microglia themselves are the primary producers of C1q within the CNS, raising the possibility of C1q can act as an autocrine mechanism for regulating microglial state. Silverman et al reports that microglial activation is completely ablated in C1q KO mice following retinal ischemia and reperfusion injury. Autocrine feedback loops are often utilized by immune cells to self-regulate state, but whether C1q could be acting in this manner is unknown and will be investigated in future studies. In parallel, the critical role of sex differences in microglia has become increasingly recognized (Lynch, 2022; Sullivan & Ciernia, 2022). The novel organelle phenotyping approach established here provides the field with a vital tool that enables investigation of autocrine mechanisms and sex differences in microglia.

## Materials and Methods

### Acquisition and maintenance of iPSC

UCI Alzheimer’s Disease Research Center (ADRC)76 cell line was obtained from human fibroblasts to generate iPSC. The cell line was derived from subject 76 (male); reprogramming and differentiation was approved by the University of California, Irvine Institutional Review Board (IRB Protocol #2013-9561). iPSC use and differentiation towards microglia was approved by the University of California, Irvine Human Stem Cell Research Oversight Committee (hSCRO protocol # 2006-5295).

iPSC were plated onto 6-well plates (Corning cat#08-772-1B) coated with growth factor-reduced Matrigel (1 mg/mL; BD Biosciences, cat#356231). iPSC cell maintenance involves daily media changes in mTeSR Plus (Stem Cell Technologies, cat#100-0276) and a humidified incubator (5% CO2, 37 C). Medium was supplemented with 0.5uM Thiazovivin (StemCell Technologies, cat#72252) for the first 24 h post-passage to promote colony survival. iPSC were tested for mycoplasma every three months and were confirmed to be negative.

### Differentiation of microglia from iPSC

iPSC were differentiated into microglia using a previously published protocol (McQuade et al., 2018). Briefly, iPSC were passaged in a 6 well-plate with mTeSR Plus at a density of 40-80 small colonies (<100 cells) per well. iPSC-derived hematopoietic progenitor cell differentiations were completed with STEMdiff Hematopoietic Kit (Stem Cell Technologies, cat#5310) where cells are fed with Media A (day 0-2) and Media B (day 3-10). Non-adherent CD43+ iHPC are collected on days 11/12 and plated in iMGL medium (Table 1), freshly supplemented with 100ng/mL IL-34 (Peprotech, cat#200-34), 50ng/mL TGFb1 (Peprotech, cat#100-21), and 25 ng/mL M-CSF (Peprotech, cat#300-25) for 28 days. The last 3 days in culture involve feeding in iMGL maturation media, which is iMGL medium freshly supplemented with 100ng/mL CD200 (Novoprotein, cat#BP004) and 100 ng/mL CX3CL1 (Peprotech, cat#300-18). Mature iMGL were treated with LPS (ThermoFisher Scientific, cat#00-4976-03), IL-10 (R&D Systems, cat#21-7IL0-10), or C1q (My Biosource, cat#MBS147305) as described.

| Product | Manufacturer |
| --- | --- |
| DMEM/F12 | Gibco, #1103901 |
| Insulin-Transferin-Selenite | Gibco, #41400045 |
| 2x B27 | Gibco, #17504044 |
| 0.5x N2 | Gibco, #17502048 |
| 1x Glutamax | Gibco, #35050061 |
| 1x NEAA | Gibco, #11140050 |
| 400 mM monothioglycerol | Sigma, #M6145 |
| 5 ug/mL human insulin | Sigma, #I2643 |

### Microglia organelle phenotyping

Following 24 hour treatment, iMGL were processed and data was analyzed as previously described (Scipioni, 2023). Briefly, cells were labeled with the following environment-sensitive dyes to observe changes in organelles: Rhodamine123 (mitochondrial membrane potential), lysotracker green (lysosomal pH), nile red (lipid droplet polarity), and hoechst (DNA). Live-cell data was acquired using LSM880.

### qPCR

Following treatment, cells were lysed using RLT buffer mastermix (10 uL beta-mercaptoethanol : 1 mL RLT) and cells were lysed with RNAeasy micro kit (Qiagen, 74004). RNA quality and quantity was validated using a nanodrop (ThermoFisher Scientific). Any remaining genomic DNA from the collected samples was degraded using RNA clean up kit (Invitrogen, cat#:AM1906). RNA samples were th converted to cDNA using high-Capacity RNA-to-cDNA kit (Applied Biosystems, cat#:4387400). qPCR reaction was performed in a 10 uL volume using the qPCR primers listed in Table 2 (ThermoFisher Scientific, cat#: 4448892). Reaction included 20x TaqMan Gene Expression Primer, 2x TaqMan Gene Expression Fast Mastermix (Applied Biosystems, cat#:4444557), cDNA template, and RNAse free water. Plate was sealed, centrifuged briefly, and loaded into Quantstudio7. Data was analyzed using the comparative delta-delta Ct method.

| Gene | Assay ID |
| --- | --- |
| 18s | Hs03003631_g1 |
| CCL2 | Hs01574247_m1 |
| IL1b | Hs01555410_m1 |
| IL6 | Hs00174131_m1 |
| IL-10 | Hs00961622_m1 |
| BDNF | Hs02718934_s1 |
| IGF1 | Hs01547656_m1 |
| P2RY12 | Hs01881698_s1 |
| SELPLG | Hs05033974_s1 |
| CX3CR1 | Hs01922583_s1 |

### Transwell Migration Assay

iMGL were collected and resuspended to a concentration of 300,000 cells/mL. 30,000 cells (100 uL) of the single-cell resuspension was added to each migration assay chamber, before placing chambers into feeder trays (Millipore) containing 150 uL of media for each condition; the chambers were then incubated at 37 °C for 3 hr. Subsequently, the chambers were transferred onto new 96-well trays containing 150 mL of prewarmed cell detachment buffer and incubated for 30 min at 37 °C. At the end of this incubation, 50 mL 1:75 dilution of CyQuant GR Dye:Lysis buffer was added to the cell detachment buffer and incubated for 15 min at room temperature. Finally, 150 mL CyQuant GR Dye:Lysis/detachment solution was transferred to a new 96-well plate, and migration was quantified using a 480/520 nm filter set on a fluorescent plate reader. All experiments were conducted in biological triplicate with technical triplicates.

### Flow cytometry (Click-it EdU and AnnexinV)

All samples were acquired on the BDS Fortessa and data was analyzed using FloJo. All experiments were conducted in biological triplicate and technical triplicate. <u>EdU</u>: Cell proliferation was assessed using Click-iT EdU flow cytometry assay kits (Life technologies, cat#: C10419). iMGL were incubated with EdU 1:000 for 3 hours prior to treatment completion in the incubator. iMGL were collected and fixed with 4% PFA for 15 minutes. Then, cells were washed in PBS-1%FBS and permeabilized with 0.1% saponin in PBS-1%FBS for 15 minutes. Lastly, cells were incubated with staining solution [1x click-it reaction buffer, CuSO4, AF488 picolyl azide, and 1x additive] for 45 minutes at room temperature and washed with PBS-1% FBS before acquisition. <u>Annexin</u>: Apoptossi was assessed using PE Annexin V Apoptosis Detection Kit with 7-AAD (Biolegend, cat#: 640934). iMGL were collected and washed twice with Biolegend’s cell staining buffer. Cells were resuspended in 100 uL Annexin V binding buffer and treated with PE Annexin V 1:20 and 7-AAD viability staining solution 1:20. Antibodies were incubated for 15 minutes room temperature and 400 uL of Annexin C binding buffer was added to each tube before acquisition.

### Apolive Glo Cell Viability/Apoptosis Assay

iMGL viability and apoptosis was quantified with ApoLive-Glo™ Multiplex Assay (Promega, cat#: G6410) per manufacturer’s instructions. Briefly, iMGL were treated with GF-AFC, a cell-permeant substrate that becomes fluorescent when it enters the cell to quantify viable cells. Then, cells are lysed and treated with a luminogenic caspase-3/7 substrate to quantify amount of caspase present. Data is shown as caspase luminescence/live cell fluorescence as a normalization measure. All experiments were conducted in n=3 biological replicates and n=5 technical replicates.

